# Fast automated reconstruction of genome-scale metabolic models for microbial species and communities

**DOI:** 10.1101/223198

**Authors:** Daniel Machado, Sergej Andrejev, Melanie Tramontano, Kiran Raosaheb Patil

## Abstract

Genome-scale metabolic models are instrumental in uncovering operating principles of cellular metabolism and model-guided re-engineering. Recent applications of metabolic models have also demonstrated their usefulness in unraveling cross-feeding within microbial communities. Yet, the application of genome-scale models, especially to microbial communities, is lagging far behind the availability of sequenced genomes. This is largely due to the time-consuming steps of manual curation required to obtain good quality models and thus physiologically meaningful simulation results. Here, we present an automated tool – CarveMe – for reconstruction of species and community level metabolic models. We introduce the concept of a universal model, which is manually curated and simulation-ready. Starting with this universal model and annotated genome sequences, CarveMe uses a top-down approach to build single-species and community models in a fast and scalable manner. We build reconstructions for two model organisms, *Escherichia coli* and *Bacillus subtillis*, as well as a collection of human gut bacteria, and show that CarveMe models perform similarly to manually curated models in reproducing experimental phenotypes. Finally, we demonstrate the scalability of CarveMe through reconstructing 5587 bacterial models. Overall, CarveMe provides an open-source and user-friendly tool towards broadening the use of metabolic modeling in studying microbial species and communities.

## Introduction

Linking metabolic phenotype of an organism to environmental and genetic perturbations is central to several basic and applied research questions. To this end, genome-scale metabolic models provide a mechanistic basis allowing to predict the effects of, e.g., gene knockouts, or nutritional changes [1, 2] Indeed, such models are currently used in a wide range of applications, including rational strain design for industrial biochemical production [3, 4], drug discovery for pathogenic microbes [5], and the study of diseases with associated metabolic traits [6, 7, 8, 9].

An emerging application of genome-scale models is the study of cross-feeding and nutrient competition in microbial communities [10, 11, 12, 13, 14, 15]. However, a vast majority of relevant microbial communities, such as those residing in the human microbiota [16], ocean [17] or soil [18], still remain inaccessible for metabolic modeling due to the unavailability of the corresponding species-level models. Thus, applications of metabolic modeling lag behind the opportunities presented by the increasing number of genomics and metagenomics datasets [19]. A major bottleneck is the so-called genome-scale reconstruction process, which often requires laborious and time-consuming curation, without which the model quality remains low. This becomes an even more stringent bottleneck considering that microbial communities can contain hundreds of different species.

Several metabolic reconstruction tools are currently available, each offering different degrees of trade-off between automation and human intervention [20, 21, 22, 23]. These tools follow a bottom-up reconstruction approach consisting of the following main steps: 1) annotate genes with metabolic functions; 2) retrieve the respective biochemical reactions from a reaction database, such as KEGG [24]; 3) assemble a draft metabolic network; 4) manually curate the draft model. The last step includes several tasks, such as adding missing reactions required to generate biomass precursors (gap-filling), correcting elemental balance and directionality of reactions, detecting futile cycles, and removing blocked reactions and dead-end metabolites (see [25] for a detailed protocol). If these problems are not resolved, the model can generate unrealistic phenotype predictions, such as incorrect biomass yields, excessive ATP generation, false gene essentiality, or incorrect nutritional requirements. The manual curation step is time-consuming and includes repetitive tasks that must be performed for every new reconstruction.

In this work, we present CarveMe, a new reconstruction tool that shifts this paradigm by implementing a top-down reconstruction approach (Fig. 1). We begin by reconstructing a universal metabolic model, which is manually curated for the common problems mentioned above. Notably, this universal model is simulation-ready: it includes import/export reactions, a universal biomass equation, and contains no blocked or unbalanced reactions. Following, for every new reconstruction, the universal model is converted to an organism-specific model using a process called “carving” (see Methods section for details). In essence, this process removes reactions and metabolites not predicted to be present in the given organism, while preserving all the manual curation and relevant structural properties of the original model. The lack of manual intervention makes this process easily automatable and the reconstruction of multiple species can be easily parallelized (see legend Fig. 1). CarveMe can build models from genome sequences alone, but allows users to optionally provide collected experimental data (in a simple tabular format) as additional input during reconstruction. Furthermore, it automates the creation of microbial community models by merging selected sets of single-species models into community-scale networks.

**Figure 1.**
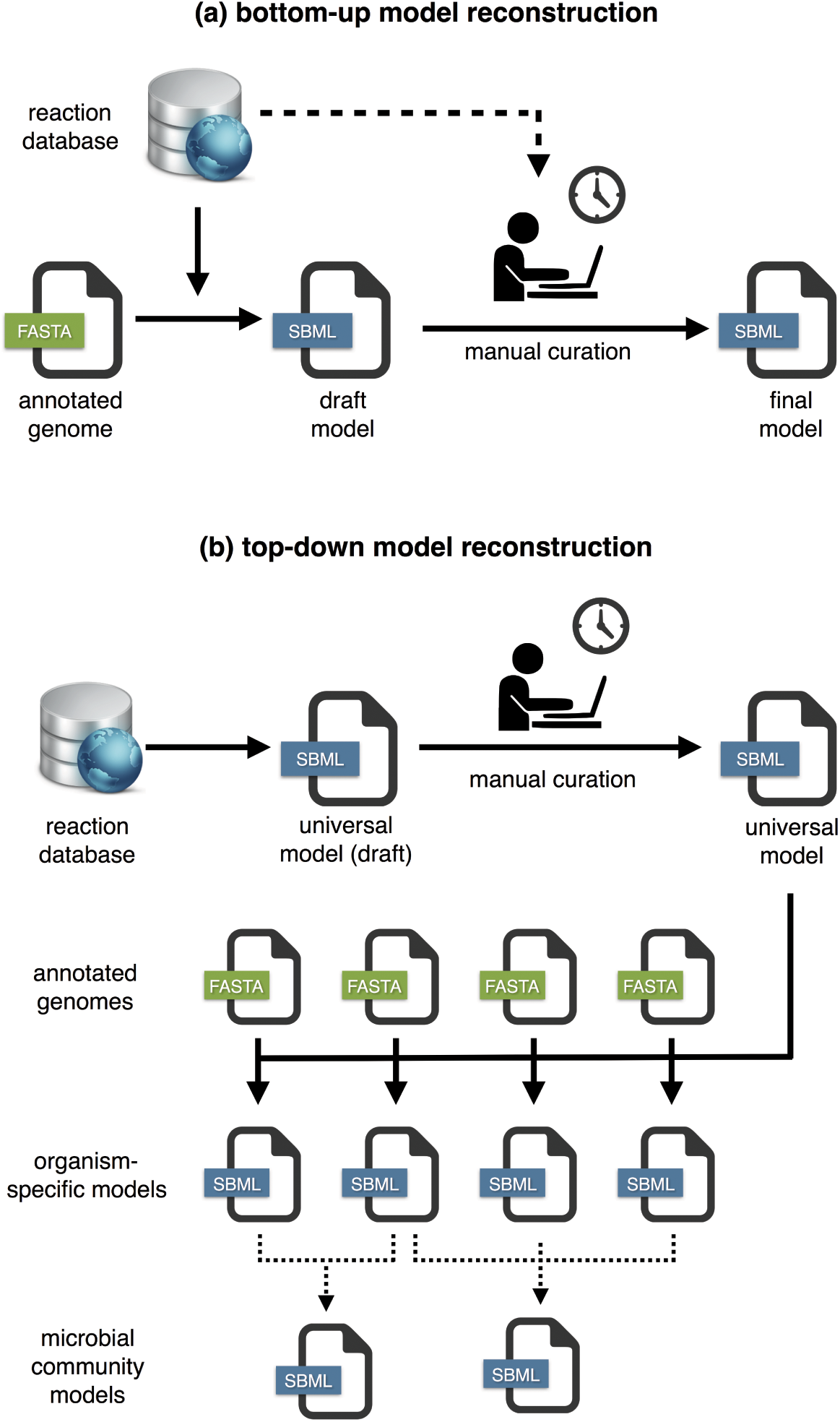
Workflow comparison between bottom-up and top-down genome-scale metabolic model reconstruction: a) In the traditional (bottom-up) approach, a draft model is automatically generated from the genome of a given organism, followed by extensive manual curation; b) In the top-down approach, a universal model is automatically generated and manually curated. This model is then used as a template for organism-specific model generation (*carving*), a process that does not require manual intervention and can be easily parallelizable to automatically generate a large number of models. Optionally, this automated process can be also applied to the generation of microbial community models by merging single-species models.

## Results

### Universal model of bacterial metabolism

We built an universal model of bacterial metabolism by downloading reaction data from BiGG [36], a database that integrates data from 79 genome-scale metabolic reconstructions (accessed: version 1.3). Although BiGG includes a few eukaryotic reconstructions (such as yeast, mouse, and human), prokaryotes, especially bacteria, are better represented, including species from 14 different genera. The universal model contains all BiGG reactions with the exception of those exclusive to eukaryotic organisms. The model underwent several curation steps (see Methods for details), including estimation of thermodynamics, verification of elemental balance, elimination of energy-generating cycles, and integration of a core biomass composition. The final model includes 3 compartments (cytosol, periplasm, and extracellular), 2383 metabolites (representing 1503 unique compounds) and 4383 reactions (2463 enzymatic reactions, 1387 transporters, and 473 metabolite exchanges). From this universal model, we derived two specialized templates for gram-positive and gram-negative bacteria with modified biomass equations to account for the respective differences in cell wall/membrane composition. During reconstruction the user can select between the generic or specialized (or even user-provided) universal models. In addition to the model, we generate a sequence database with a total of 30814 unique protein sequences derived from the gene-protein-reaction associations of the original models. These are used to align the input genomes and identify the respective reactions in the universal model.

### Comparison with manually curated metabolic models: Escherichia coli and Bacillus subtilis

The quality of the models generated with CarveMe was evaluated regarding their ability to reproduce experimental data and compared to previously published manually curated models. Two reference model organisms were selected as case-studies, namely the gram-negative bacterium *Escherichia coli* (strain K-12 MG1655) and the gram-positive bacterium *Bacillus subtilis* (strain 168). *E. coli* is among the best-studied organisms and the one with the most highly curated metabolic models. *B. subtilis* was chosen as a representative of gram-positive bacteria due to the availability of experimental data to test model predictions. The genomes of these two species used to build the models were downloaded from NCBI RefSeq (release 84) [26]. For comparison, the manually curated models for *E. coli* (iJO1366) and *B. subtilis* (iYO8444) were obtained from the respective publications [27, 28] Furthermore, to analyze how CarveMe performs in comparison to the currently available automated reconstruction tools, the same genomes were used to generate genome-scale models with the modelSEED pipeline (https://kbase.us/ accessed: August 2017) [20]. Notably, the models generated with CarveMe were able to reproduce growth on M9 minimal medium from genome data alone, without specifying any growth requirements during reconstruction or performing any additional gap-filling after reconstruction. The models generated with modelSEED required specific gap-filling for M9 minimal medium.

All models were next used to simulate growth on Biolog phenotype arrays (see Methods) and their predictive ability was assessed with respect to the experimental data in terms of accuracy, precision, sensitivity, specificity, and F1-score. It can be observed that the CarveMe *E. coli* reconstruction performs very similarly to the iJO1366 model with regard to all the performance metrics considered, with the exception of slightly decreased specificity (Fig. 2a). The modelSEED *E. coli* model shows higher specificity but lower performance with regard to the other metrics.

**Figure 2.**
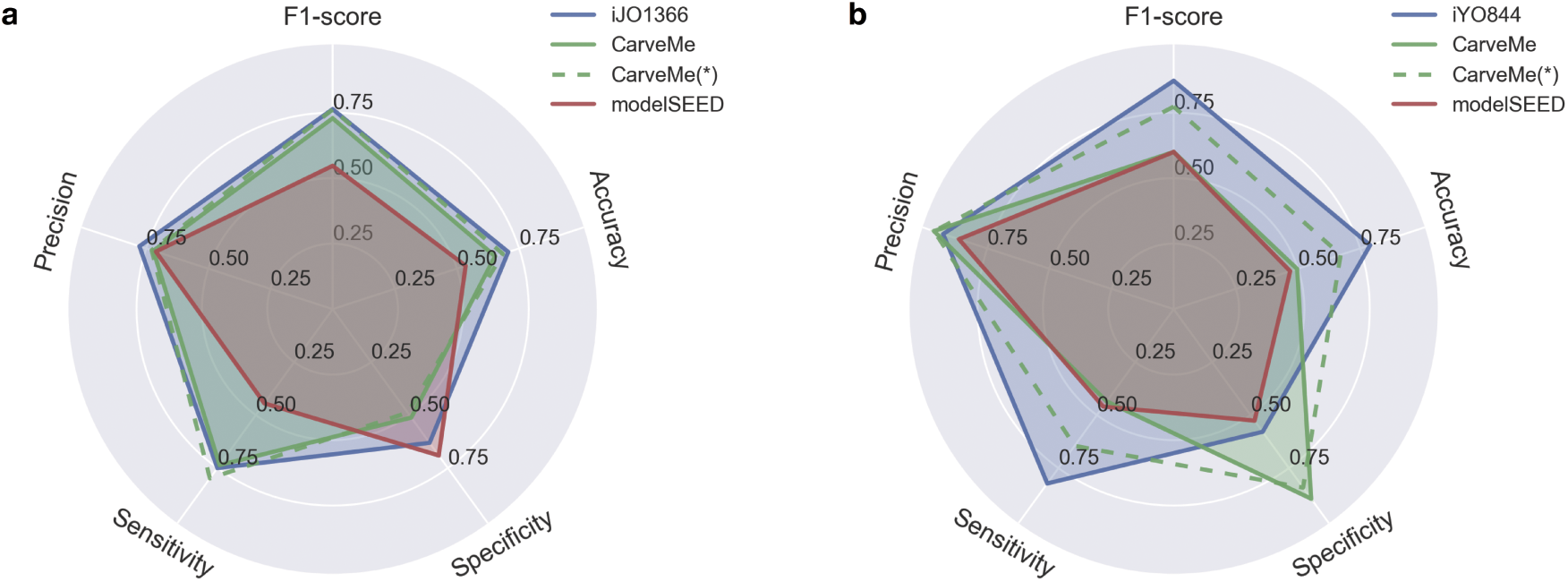
Phenotype array simulation results: a) *E. coli* models: CarveMe reconstruction vs modelSEED reconstruction vs iJO1366 model; b) *B. subtilis* models: CarveMe reconstruction vs modelSEED reconstruction vs iYO844 model. The dotted lines represent the simulation results for CarveMe models when free diffusion is allowed for all metabolites without associated transporters.

In the case of *B. subtilis*, the CarveMe model performs with high specificity compared to the iYO844 model but looses on other metrics (Fig. 2b). The modelSEED model performs worse than iYO844 with regard to all the metrics. This difference in performance with respect to the iYO844 model can be explained by the lack of annotated transporters for *B. subtilis*, which have been manually curated in this model. To test this hypothesis, we repeated all simulations with the CarveMe models, this time allowing free diffusion of metabolites without associated transporters (Fig. 2, dotted lines). It can be observed that, for *B. subtilis*, the model performs better with regard to all metrics, whereas for *E. coli* the results remain practically unaltered.

The reconstructed models were then evaluated for their ability to predict gene essentiality (see Methods). In the case of *E. coli*, all models perform similarly (Fig. 3a). On the other hand, the automated reconstructions for *B. subtilis* performed with lower precision and sensitivity compared to the iYO844 model (Fig. 3b). One common pattern for both CarveMe models is a slightly decreased sensitivity with regard to gene essentiality (i.e. many essential genes reported as false negatives). One possible reason is the utilization of generic biomass templates that exclude non-universal cofactors [29] and other organism-specific biomass precursors. To test this, we generated new reconstructions using the biomass composition present in the manually curated models and repeated the simulations (Fig. 3, dotted lines). Confirming to our hypothesis, we observed a gain in sensitivity when using organism-specific biomass composition.

**Figure 3.**
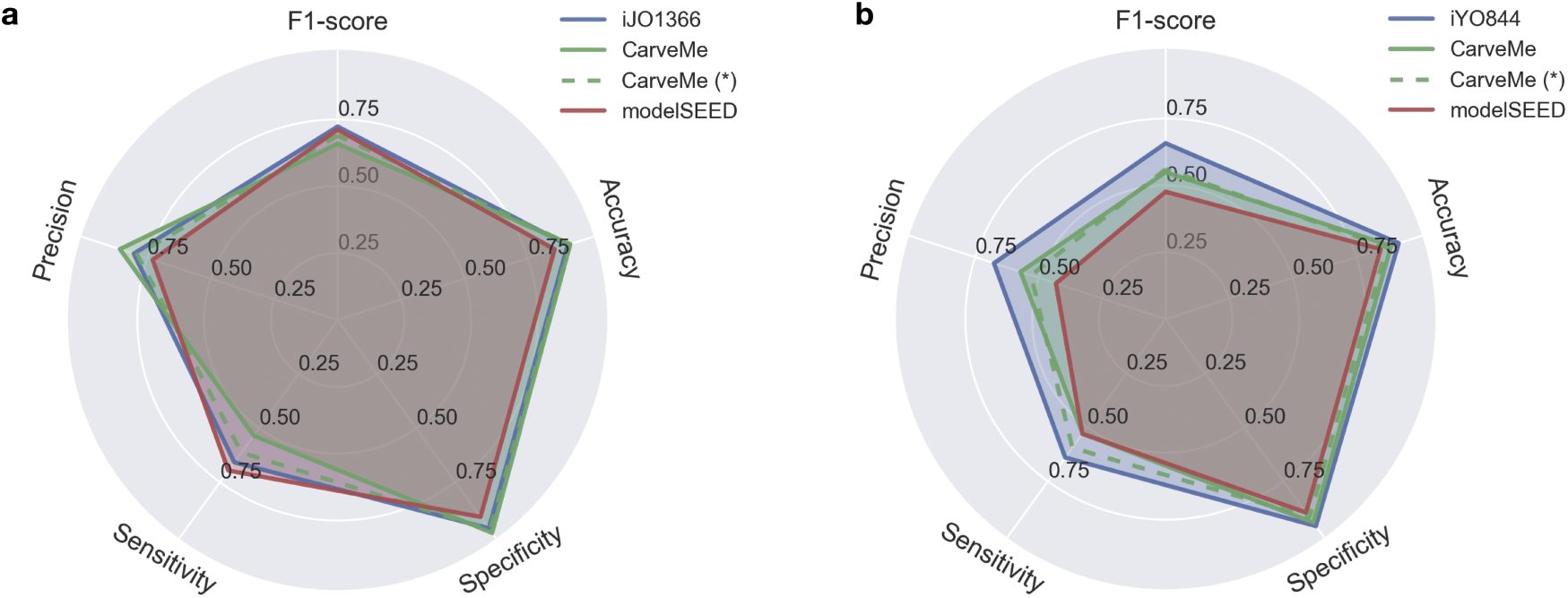
Gene essentiality prediction results: a) Gene essentiality for *E. coli* growing in minimal medium using glucose or glycerol as carbon source (combined results); b) Gene essentiality for *B. subtilis* growing in rich medium. The dotted lines represent the simulation results for CarveMe models when the original biomass composition of the respective organism is used during reconstruction.

A general limitation in building genome-scale metabolic models, given the available data, is the existence of several equally plausible models. This has been explored by [30], where the authors show that when gap-filling a model for multiple growth media, the selected order of the media influences the final network structure. Another source of ambiguity is the degeneracy of solutions when solving an optimization problem to generate the model itself. To tackle this issue, CarveMe allows the generation of model ensembles. The user can select the desired ensemble size, and then use the ensemble model for simulation purposes in order to explore alternative solutions. We generated model ensembles (N=100) for *E. coli* and *B. subtilis* and repeated the phenotype array simulations and gene essentiality predictions using different voting thresholds (10%, 50%, 90%) to determine a positive prediction (see Methods). To confirm that the ensemble generation is unbiased, we calculated the Jaccard distance between every pair of models in each ensemble (Supp. Fig. 1a). The ensemble models show a degree of variability with the pairwise distance following a normal distribution. The ensemble for *B. subtilis* shows a much larger variability compared to *E. coli*, reflecting a higher degree of uncertainty in the network structure. Regarding the phenotype array simulations, we observed a general decrease in the specificity for both *E. coli* and *B. subtilis* (Supp. Fig. 1b,c). The difference is more evident for *B. subtilis*, simultaneously displaying a considerable increase in the other metrics. Interestingly, the predictions seem to be consensual irrespective of the voting threshold applied. This is also evident for the gene essentiality predictions, where the results are unaltered with regard to the single model simulations (Supp. Fig. 1d,e). One exception is the *B. subtilis* ensemble for voting thresholds above 90%, where no positive consensus is ever obtained.

Taken together, these comparisons demonstrate that, considering the fast and fully automated procedure, the CarveMe reconstructed models perform on close level to the manually curated models.

### Metabolic models for the human gut bacterial species

The human gut microbiome is of particular interest for metabolic modeling due to its impact on human health [31, 32, 33]. In this regard, one major hurdle is the characterization of gut bacterial species in terms of growth requirements and metabolic potential. In a recent study from our group, the growth of 96 phylogenetically diverse gut bacteria was characterized across 19 different media (15 defined and 4 complex media compositions) (Tramontano *et al, in revision*). From this list, a total of 47 bacteria are also included in the AGORA collection, a recently published collection of 773 semi-manually curated models of human gut bacteria [34]. These models, however, could not reproduce the metabolic needs experimentally observed by Tramontano et al., highlighting the need for the use of validated growth media for species that are distant from model organisms.

In this work, we used CarveMe to reconstruct the metabolism of 74 bacterial strains that grew in at least one defined medium in the Tramontano et al. study. We additionally used a recently published collection of literature-curated uptake/secretion data for human gut species to further refine the reconstructed models [35]. The resulting models thus represent an up-to-date and experimentally backed collection for gut bacteria. To understand how the media information contributed to the model refinement, we analysed the gap-filling reactions introduced in each model when these data is provided during reconstruction. Interestingly, we observed that the total number of gap-filling reactions is negatively correlated with the genome size of the respective organism (Pearson’s r = -0.49, p < 10^−5^; Supp Fig. 2a). This is most likely due to poor annotations of genes in amino acid metabolism, especially transporters, that are responsible for several of the gap-filled pathways in models of small-genome species (which usually tend to harbor many auxotrophies). We next compared the number of gap-filled reactions between CarveMe and AGORA models (40 models in common) and observed that they are well correlated (Pearson’s r = 0.52, p < 10^−3^; Supp. Fig. 2b). This shows that the quality of the CarveMe models even without the Tramontano *et al* data is comparable to the manually-curated AGORA collection, which further highlights the advantages of our top-down curation approach.

### Large-scale model reconstruction and community models

To demonstrate the scalability of CarveMe, and to provide the research community with a collection of reconstructed models spanning a wide variety of microbes, we reconstructed 5587 bacterial genome-scale models. These correspond to all bacterial genomes available in NCBI RefSeq (release 84) [26] that were classified as reference or representative assemblies at the strain level. This collection of models represents metabolism across all (currently sequenced) bacterial life and thus can help uncovering principles underlying the architecture and diversity of metabolic networks.

The smaller metabolic networks belong to the Mycoplasma genus with as few as 238 reactions (*Mycoplasma ovis* str Michigan), whereas the larger metabolic networks belong to the *Klebsiella* and *Escherichia* genera, with up to 2472 reactions (*Klebsiella oxytoca* str CAV1374). We note that these numbers are indicative, as they can be biased or restricted by the quality of the gene annotation and the scope of the reaction databases. Interestingly, the total number of reactions and metabolites per organism appears to vary by approximately one order of magnitude across all bacteria (Fig. 4a). Furthermore, the number of metabolites per species seems to scale linearly with the number of reactions (average metabolite/reaction ratio of 0.6), albeit with an asymptotic trend closer to the maximum number of reactions. The latter indicates saturation of metabolite space relative to the expansion of the reaction space due to, for example, alternative pathways acting on the same metabolites. This pattern, however, does not completely account for the secondary metabolism, since it yet remains largely unknown.

**Figure 4.**
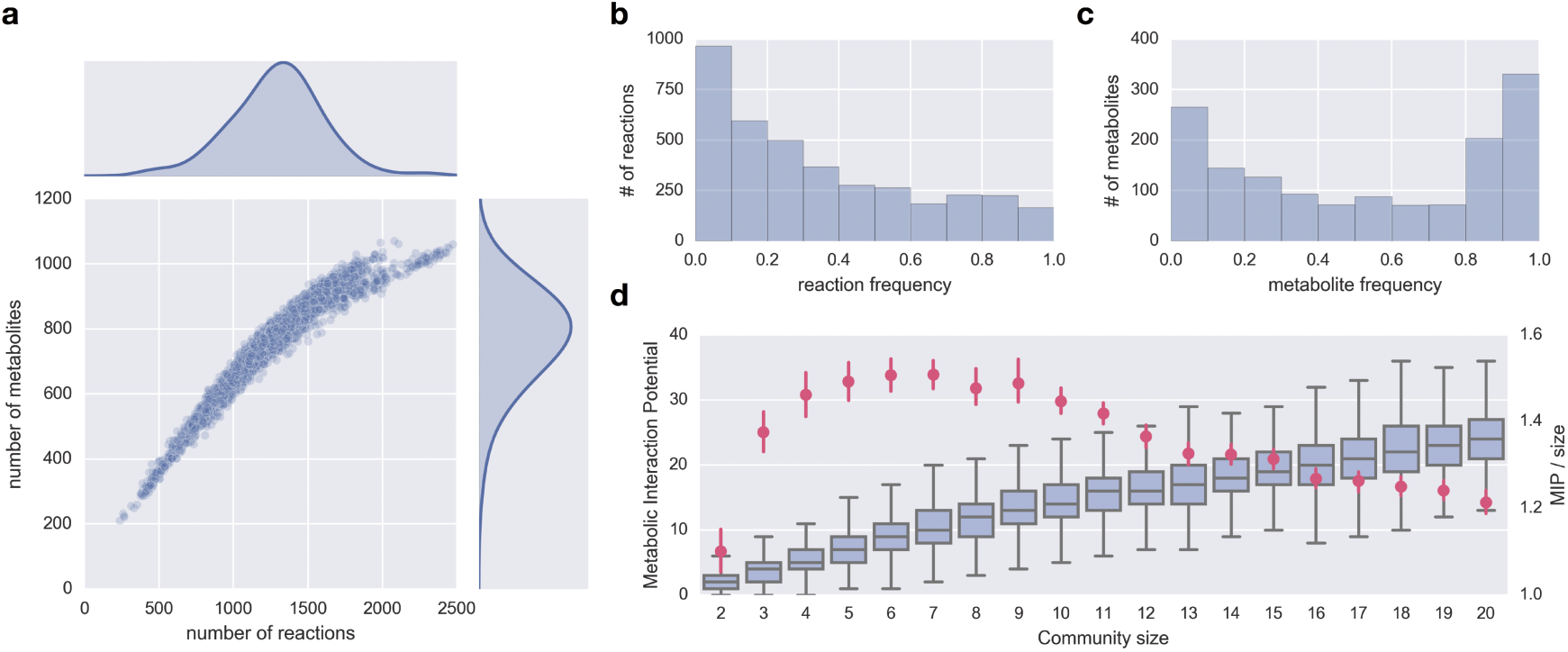
NCBI RefSeq reconstruction collection summary: a) Total number of reactions and metabolites per organism; b) Reaction frequency distribution across all organisms; c) Metabolite frequency distributions across all organisms; d) Metabolic interaction potential of randomly assembled communities of different sizes, including absolute MIP values (box plots), and average MIP normalized by community size (red dots with 95% confidence intervals).

We further looked at how frequently individual reactions and metabolites occur across species. The frequency of reactions shows a negative exponential distribution (Fig. 4b) with 30% of all reactions being present in less than 10% of the organisms, and only 5% of reactions being present more than 90% of the organisms. Interestingly, the metabolite frequency shows instead a bimodal distribution (Fig. 4c) with peaks below 10% frequency (≈ 20% metabolites) and above 90% frequency (≈ 25% metabolites). As expected, the high frequency reactions and metabolites are distributed across the primary metabolism, including carbohydrate, lipid, nucleotide, amino acid and energy metabolism (Supp. Fig. 3).

Next, we illustrate the use of our model collection to build microbial community models and explore inter-species interactions. We created random assemblies of microbial communities of different sizes (up to 20 member species per community, 1000 assemblies for each community size) and calculated their metabolic interaction potential (MIP score, as defined in [11]). In brief, the MIP score of a community is a measure of the number of compounds that can be exchanged between the community members, allowing the community to reduce its dependence on the environmental supply of nutrients. The results obtained with this collection (Fig. 4d) closely match those previously obtained with a smaller model collection (1503 bacterial strains, generated with modelSEED) [11]. In particular, while the potential for interaction increases with the community size, the number of interactions normalized by the community size shows a maximum at a relatively small community size.

## Discussion

CarveMe uses the BiGG database [36] to build universal model templates, which offers some advantages compared to other commonly used reaction databases such as KEGG, Rhea, or modelSEED [24, 20, 37]. First, it was built by merging several genome-scale reconstructions (ranging from bacterial to human models), which leverages on the curation efforts that were applied to these models. Secondly, BiGG reactions contain gene-protein-reaction associations, compartment assignments, and human-readable identifiers. These aspects facilitate the reconstruction process and the utilization of the generated models. However, BiGG is limited in size and scope compared to other databases, which may result in lack of coverage of certain metabolic functions, especially for secondary metabolism. A comparison between the reaction content of BiGG, modelSEED and KEGG (Supp. Fig. 4), shows that BiGG contains a large number of reactions which are not present in the other databases. However, when comparing the total coverage of unique EC numbers, one can observe that BiGG lags behind modelSEED and KEGG. Furthermore, transport reactions are often missing from these databases due to the lack of functional annotation of transporter genes [38], which can hamper the correct prediction of metabolic exchanges. In future releases, we plan to facilitate the integration of reaction data from multiple databases.

The quality of the universal model used for top-down reconstruction is of paramount importance. There is a tradeoff between the desired broadness of this “universal” model and the amount of curation effort required. In this work, we opted to provide a well-curated universal model of bacterial metabolism, which can be generally used for reconstruction of any bacterial species. We also provide more refined gram-positive and gram-negative template models, which account for the differences in membrane/cell wall composition. The models generated with CarveMe are pre-curated for common structural and biochemical inconsistencies and can be readily used for simulation. Nonetheless, they should still be considered as drafts subject to further refinement, as they might require organism-specific curation to reproduce certain phenotypes. In future releases, we aim to provide a larger collection of templates in order to improve coverage of all domains of microbial life. This includes other prokaryotes (archaea) and eukaryotes such as yeasts. Currently, the pipeline allows users to provide their own templates at any desired taxonomic level, and provides methods to facilitate the creation and curation of such templates (see Methods). Note that it is also possible to utilize the pipeline in a “traditional” reconstruction approach, i.e. to use the complete reaction database as a non-curated universal model, and perform the manual curation of the generated models *a posteriori*.

In conclusion, CarveMe provides an automated reconstruction solution that uses a manually curated universal model as a scaffold to generate genome-specific models. The underlying carving procedure can work with the genetic evidence alone, i.e. without requiring the specification of a medium composition. This distinguishes CarveMe from other reconstruction algorithms, and makes it especially suitable for building models for uncultured species and communities. Furthermore, when experimental data is collected for a given species (such as growth media, known auxotrophies, metabolic exchanges, presence/absence of reactions) the user can provide these data in a simple tabular format as additional input to the pipeline. We observe that the models generated for *E. coli* and *B. subtilis* remarkably reproduced growth on M9 minimal medium without media information or any other data being provided during reconstruction. Our extensive benchmarking results show that the quality of CarveMe models is highly comparable to those of the manually curated models. Another salient feature of CarveMe is its speed and easy parallelization. A single reconstruction required, on average, 3 minutes on a laptop computer (Intel Core i5 2.9 GHz). Indeed, this allowed us to rapidly create one of the largest publicly available metabolic model collections. Finally, we have given particular attention to making CarveMe an easy-to-use tool. The pipeline is available as an open-source command line tool, and can be easily installed on a personal computer as well as on a high-performance computing cluster. It implements a modular architecture that facilitates modification/integration of new components. For instance, sequence alignments can be automatically performed with diamond [42] or extracted from the output of third-party tools (we currently support eggnog-mapper [43]). Several other features such as direct download of genome sequences from NCBI RefSeq or GenBank, automatic parallelization for multiple genomes, and generation of community models, are expected to facilitate the use by a broad range of researchers.

## Methods

### Universal model building

CarveMe provides a script to build an universal draft model of metabolism by downloading all reactions and metabolites in the BiGG database [36] into a single SBML file. All metadata associated with the reactions and metabolites are stored as SBML annotations. The same script can be used to additionally perform automated curation tasks to build a final universal model (as described next). BiGG version 1.3 was used during the course of this work.

### Universal bacterial model

The initial draft model built from the entire BiGG database was manually curated to generate a functioning universal model of bacterial metabolism. In the first step, all reactions and metabolites associated with non-bacterial compartments were removed. The second step consisted on constraining reaction reversibility in order to eliminate thermodynamically infeasible phenotypes. Lower and upper bounds for the Gibbs free energy change for each reaction (Δ*G*_*r*_) were estimated using the formula:

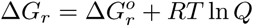

where 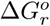 was calculated using the component contribution method [39]. R is the universal gas constant, and the temperature *T* was set to 298.15 K (25 ^o^C). The reaction quotient (Q) denotes the product to substrate ratio, which was limited by the physiological bounds imposed on metabolite concentrations. A recent study showed that absolute metabolite concentrations are conserved among different kingdoms of life [40]. The concentrations measured in this study were used as reference and allowed to vary by 10-fold with respect to the measured values. All other metabolites were allowed to vary between 0.01 and 10 mM. The reactions were set as irreversible in the thermodynamically feasible direction whenever the estimated bounds for Δ*G*_*r*_ were either strictly positive or strictly negative. Since 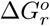 could not be determined for all reactions, additional heuristic rules were applied: 1) ATP-consuming reactions were not allowed to proceed in the reverse direction; 2) reactions present in thermodynamically curated models [41, 27] assumed the reversibility constraints adopted in these models.

In the next step, atomically unbalanced reactions were removed from the model. If not removed, these reactions can lead to spontaneous mass generation and unrealistic yields. An universal biomass equation was then added to the model. This equation was adapted from the *E. coli* biomass composition [27] in accordance to a recent study on universal biomass components in prokaryotes [29]. Subsequently, blocked reactions and dead-end metabolites were determined using flux variability analysis and removed from the model.

Finally, the model was simulated under different medium compositions, including minimal medium (M9 with glucose), and complete medium (all uptake reactions allowed to carry flux). The model was tested for biomass and ATP production. Unrealistic ATP generation was detected and the reversibility of the reactions involved was manually constrained in an iterative way until all problematic reactions operated in the most thermodynamically favorable direction.

The draft model and the final curated model are provided as part of the pipeline, as well as all the intermediate data used for model curation (Gibbs free energy estimations, reference metabolomics data, and list of manually curated reactions).

### Gram-positive and Gram-negative templates

The universal biomass composition does not contain membrane and cell wall components specific for gram-negative and gram-positive bacteria. This can result in false negative gene essentiality predictions for lipid biosynthesis pathways that generate these membrane/cell wall precursors. Therefore, we generated specialized templates for gram-positive and gram-negative bacteria by adding these components to the respective biomass composition. The gram positive template includes glycerol teichoic acids, lipoteichoic acids and a peptidoglycan unit. The gram negative template includes phosphatidylethanolamines, murein and a lipopolysaccharide unit. In both cases the final biomass composition is normalized to represent 1 gram of cell dry weight.

During reconstruction, the user can select the universal bacterial template or one of the specialized templates. Furthermore, a utility to automatically generate customized templates with user-provided biomass compositions is also included.

### Gene annotation

The amino acid sequences for all genes in the BiGG database were downloaded from NCBI and stored as a single FASTA file. This file is used to align the input genome files given by the user and find the corresponding homologous genes in the BiGG database. The user can provide the genome as a DNA or protein FASTA file. Note that raw genome files are not supported, since open reading frame (ORF) identification is outside the scope of the tool.

By default, gene/protein alignments are performed using DIAMOND [42]. Alternatively, the user can provide an alignment file externally generated with eggnog-mapper [43], which provides higher confidence for functional annotation by refining the alignments with orthology (rather than homology) predictions. However, this requires the utilization of high-performance computing resources.

### Reaction scoring

The gene alignment scores obtained in the previous step are mapped onto reaction scores using gene-protein-reaction (GPR) associations. The goal is to score the likelihood that a reaction is present in the given organism. Gene scores are first converted to protein scores by calculating the minimum score of all subunits that form a protein complex. If any subunit is missing, the protein is given a null score. Protein scores are then converted to reaction scores by summing the scores of all isozymes that can catalyze a given reaction. Customized GPR associations for the reconstructed organism are generated during this process.

The final scores are normalized to a median value of 1 and typically follow a log-normal distribution. Other enzyme-catalyzed reactions without genetic evidence are given a negative score (by default −1), and spontaneous reactions are given a neutral score.

### Model carving

Model carving is the process of converting a universal model into an organism-specific model by removing reactions and metabolites which are unlikely to be present in the given organism. This is performed by solving a mixed integer linear program (MILP) that simultaneously maximizes the presence of high-score reactions and minimizes the presence of low-score reactions while enforcing network connectivity (i.e. gapless pathways):

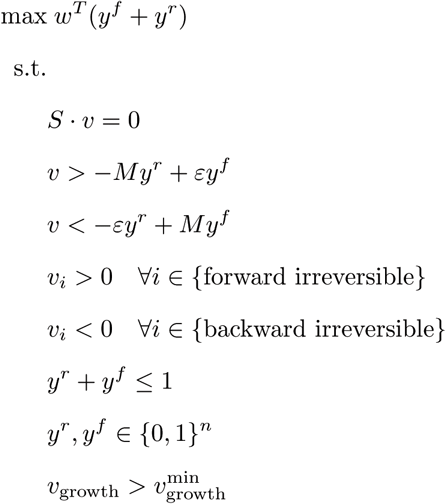

where *w* is the weighting vector (determined in the previous step), *v* is the flux vector, *y*^*ƒ*^ and *y*^*r*^ are binary vectors indicating the presence of flux in the forward or backward reaction, *S* is the stoichiometric matrix, *ε* is a minimum flux that must be carried by a reaction to be considered active (default: 0.001 mmol/gDW/h),*M* is a maximum flux value (default: 100 mmol/gDW/h), and 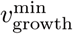 the minimum growth rate (default: 0.1 h^-1^).

After solving the MILP problem, all inactive reactions are removed from the model, including all consequently orphaned genes and metabolites. The final model is then exported as an SBML file, the standard community format (to maximize compatibility with existing tools, the user can select between the latest or the legacy version of the standard).

### Ensemble generation

CarveMe also provides the option to generate an ensemble of equally plausible models rather than a single model. This is performed by randomizing the weighting factors of reactions without genetic evidence. The MILP problem is solved multiple times with the different weighting factors to generate alternative models. The size of the ensemble (number of models) is selected by the user. The ensemble is then exported as a single SBML file.

For the purpose of phenotype array simulation and gene essentiality prediction, a voting threshold (*T*) was used to determine a positive outcome (i.e. a substrate is growth-supporting or a gene is considered essential, if the percentage of models that agree with such phenotype is larger than *T*). Simulation results with multiple thresholds (10%, 50%, 90%) are presented.

### Experimental constraints

If the user collects experimental data for an organism, these can be provided as additional input in tabular format during reconstruction and used to further constrain the optimization problem. According to the level of confidence on the collected data, these can provided as “hard” or “soft” constraints, which indicate the presence/absence and preferred direction of a given set of reactions in the model (this includes enzymatic reactions, transport reactions, and metabolite exchanges).

*Hard* constraints are simply a list of flux bounds that are applied during reconstruction 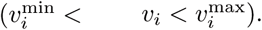. They can be used to force or block the utilization of any given reaction. (Please note that hard constraints can make the MILP problem infeasible, so they should be used with care.)

*Soft* constraints are specified as a mapping from reactions to any of three possible values (1, s−1, 0), which are used to modify the objective function, giving a different weight to the binary variables associated with the respective reactions:

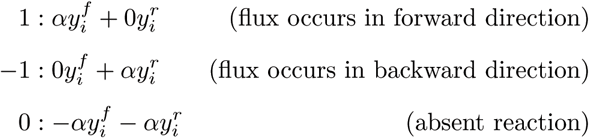

where *α* is a configurable parameter (default value: 1) that allows the user to fine-tune the priority given to the experimental evidence relatively to the genetic evidence.

### Gap-filling

CarveMe allows to gap-fill a model to reproduce growth on a set of given media. This is an optional step performed after the carving process. Its implementation is similar to typical gap-filling procedures, except that the reactions scores previously calculated are used as weighting factors. This increases the probability that a reaction with some level of genetic evidence is selected over an equivalent alternative with lower (or without) genetic evidence. The problem is formulated as follows:

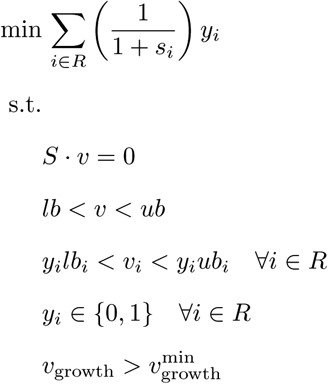

where *R* is the set of reactions in the universe not present in the model, *s*_*i*_ is the annotation score of reaction *i* (0 for reactions without score), *lb* and *ub* are the lower and upper flux bound vectors, and 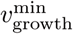 is the same as previously defined.

### Microbial community models

CarveMe provides a script to merge a selected set of single-species models into a microbial community model. The result is an SBML model where each species is assigned to its own compartment and the extracellular environment is shared between all species. There are some additional options provided, such as the creation of a common community biomass reaction, creation of isolated extracellular compartments connected by an external metabolite pool, and the initialization of the community environment with a selected growth medium. The model can be readily used for flux balance analysis (just like a single-species model) to explore the solution space of the community phenotype, and also for the application of community-specific simulation methods [10, 11]

### Simulation of phenotype arrays

Simulation of phenotype arrays was performed by constraining the respective models to M9 minimal medium (with a maximum uptake rate of 10 mmol/gDW/h for every compound). For each type of array (carbon, nitrogen, sulfur, phosphorus), the default source of the given element (respectively: glucose, ammonia, sulfate, phosphate) was iteratively replaced with the respective compounds in the array. The phenotype was considered viable if the growth rate was above 0.01 h^-1^. The experimental data used for validation was obtained from [41, 28]

### Determination of gene essentiality

Gene essentiality was determined by iteratively evaluating the impact of single gene deletions using the respective gene-protein-reaction associations. The media compositions were defined as M9 minimal medium (with glucose or glycerol as carbon source) for *E. coli* and LB medium for *B. subtilis*. The results are then validated with experimental data retrieved from [27, 28]

### Human gut bacterial species

The reconstructions were performed using the genome sequences and growth media provided in (Tramontano *et al, in revision*). The uptake/secretion data collected by [35] were provided as soft constraints during reconstruction (note that these data are reported at species, rather than strain level, hence the confidence level is low). Oxygen preferences were extracted from the PATRIC database [44] and used as hard constraints. Each model was gap-filled for the subset of media conditions where growth was observed.

### Technical details

CarveMe is implemented in Python 2.7. It requires the framed python package (version 0.4) for metabolic modeling [45], which provides an interface to common solvers (Gurobi, CPLEX), and import/export of SBML files through the libSBML API [46]. In this work, all MILP problems were solved using the IBM ILOG CPLEX Optimizer (version 12.7).

### Code Availabitity

CarveMe is publicly available with an open source license at https://github.com/cdanielmachado/carveme. It can be easily installed from a terminal by typing: pip install carveme. All the scripts used to analyse the data are available upon request during peer-review and will be publicly available upon publication.

### Data Availabitity

The database with the 5587 model collection is publicly available at https://github.com/cdanielmachado/embl_gems. The *E. coli* and *B. subtilis* models, the human gut species models, and all intermediate results are available upon request during peer-review and will be publicly available upon publication. The experimental data used to validate the different models were obtained from the supplementary materials of the respective publications as cited above.

## Acknowledgements

This project has received funding from the European Union’s Horizon 2020 research and innovation programme under grant agreement No 686070. M.T. was supported by the EMBL interdisciplinary postdoctoral program. The authors would like to thank Jaime Huerta-Cepas for implementing support for BiGG annotations in eggnog-mapper and Markus Herrgåard for critical comments on the manuscript.

## Author contributions

KP conceived the project. DM, SA and KP designed the pipeline. DM and SA wrote the code. DM performed the computational analysis. MT experimentally characterized the gut bacteria. DM and KP prepared the manuscript. All authors revised and approved the final manuscript.

## Supplementary Figures

**Supplementary Figure 1:**
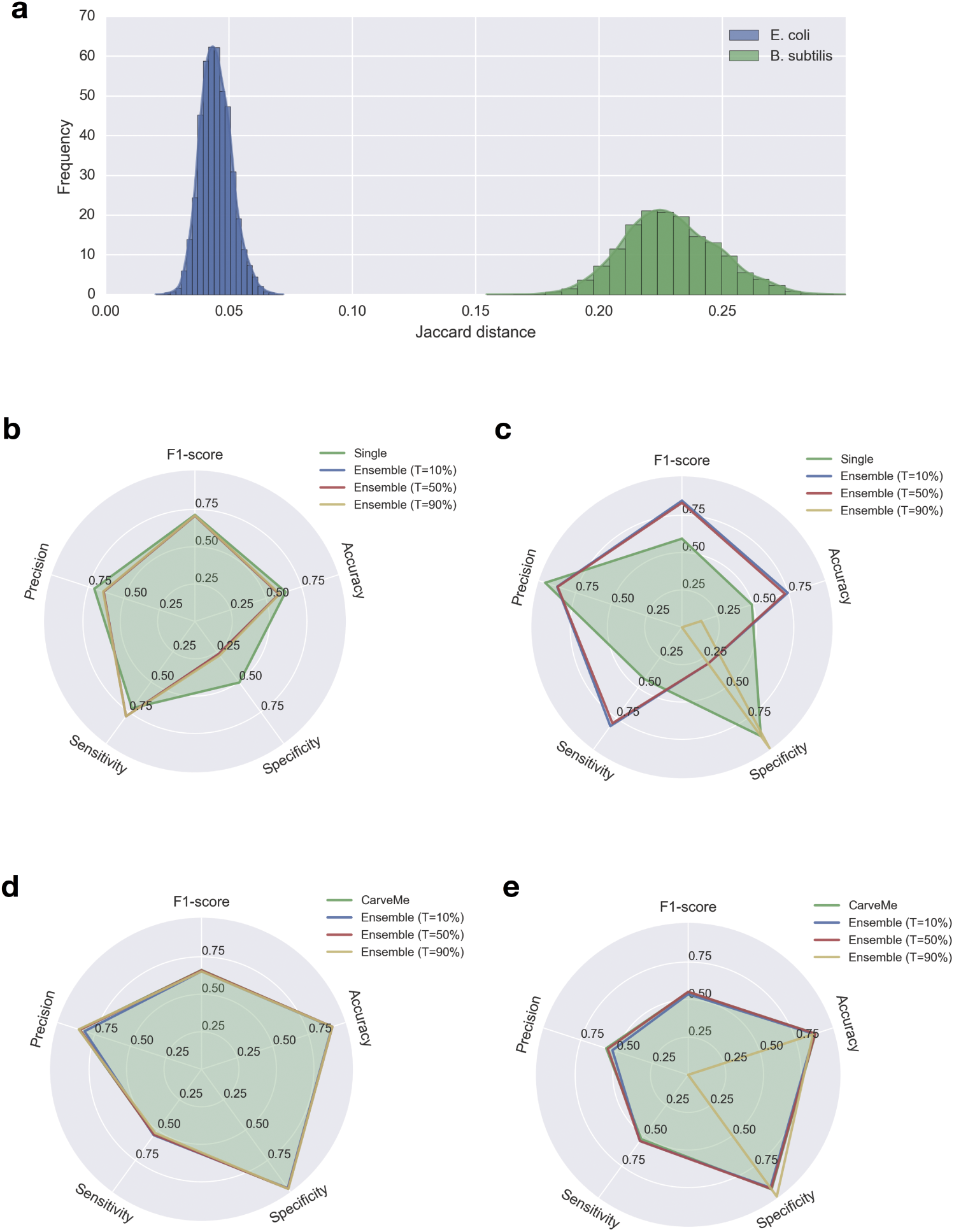
Ensemble modeling results: a) Pairwise distance between all network structures within each ensemble for the two ensembles generated (*E. coli* and *B. subtilis*); b) *E. coli* phenotype array simulation results; c) *B. subtilis* phenotype array simulation results; d) *E. coli* gene essentiality prediction; e) *B. subtilis* gene essentiality prediction. Panels b-e compare single model vs ensemble model simulation, where T denotes the voting threshold used to determine a positive result.

**Supplementary Figure 2:**
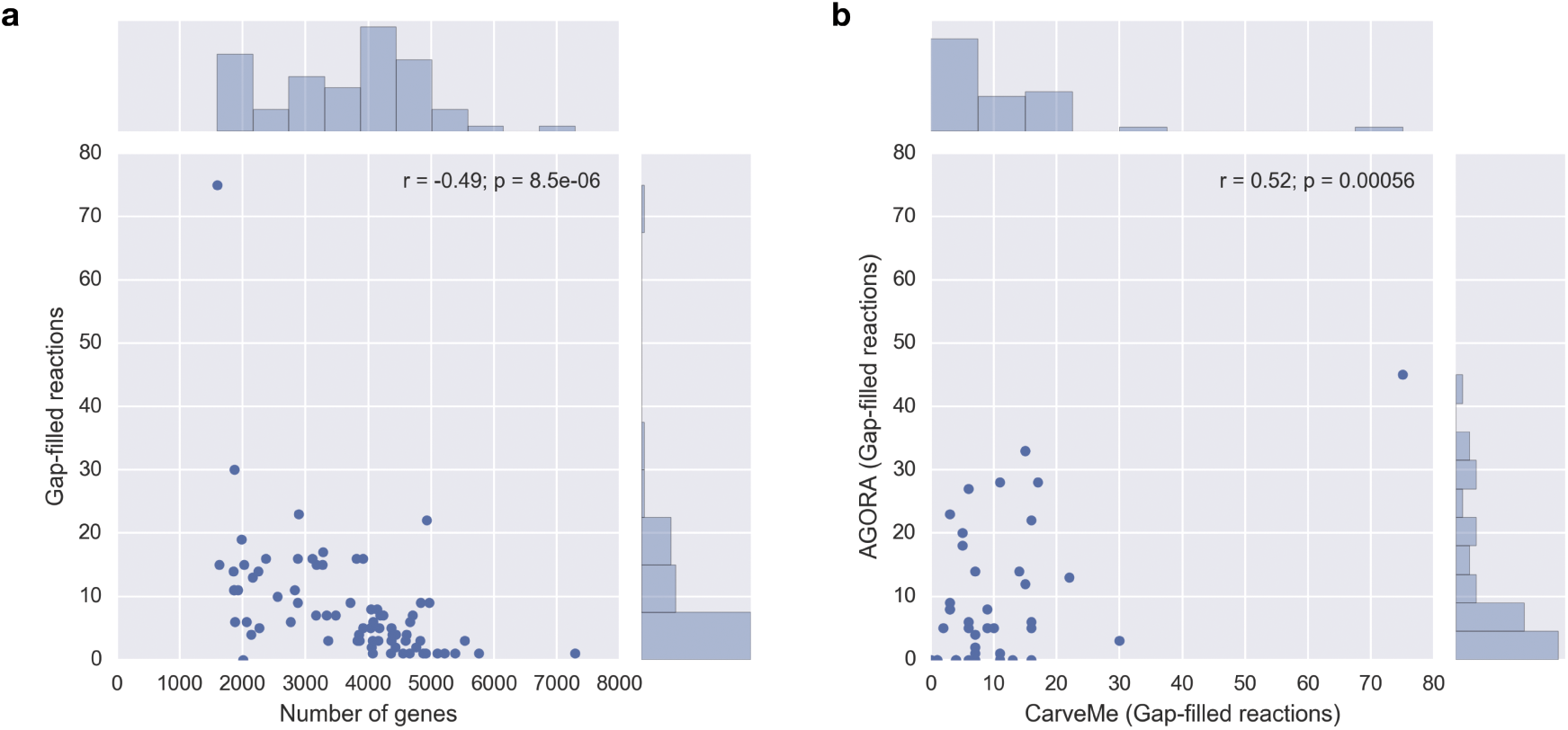
Gap-filled reactions for gut bacteria: a) Comparison between genome size and the number of gap-filled reactions per species; b) Comparison of the number of gap-filled reactions between CarveMe and AGORA models. Pearson correlation is indicated in both panels.

**Supplementary Figure 3:**
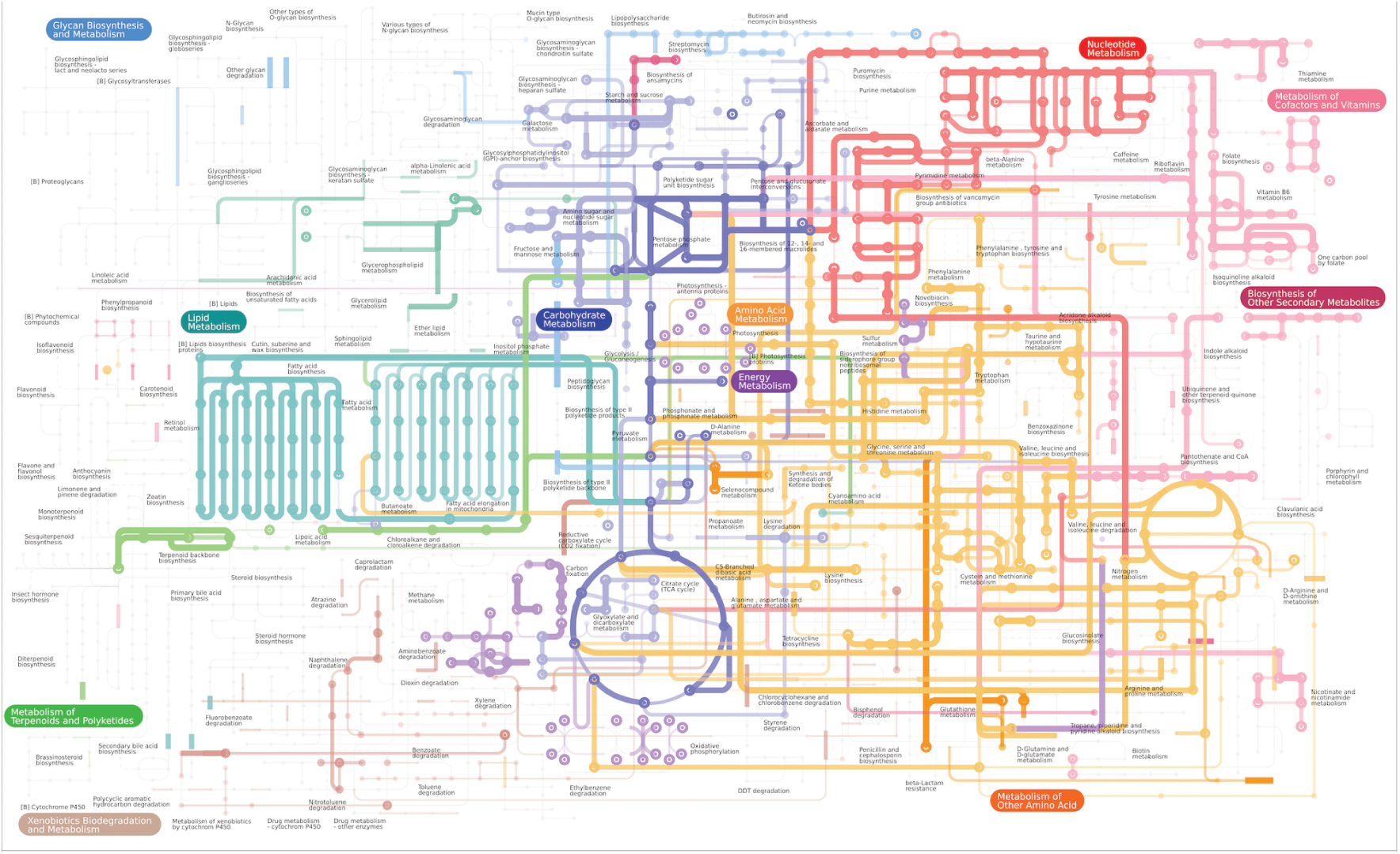
Global distribution of reaction and metabolite frequency across the bacterial model reconstruction collection. The frequency of metabolites and reactions is indicated, respectively, by node and edge width and opacity. Image generated with iPath2 [47].

**Supplementary Figure 4:**
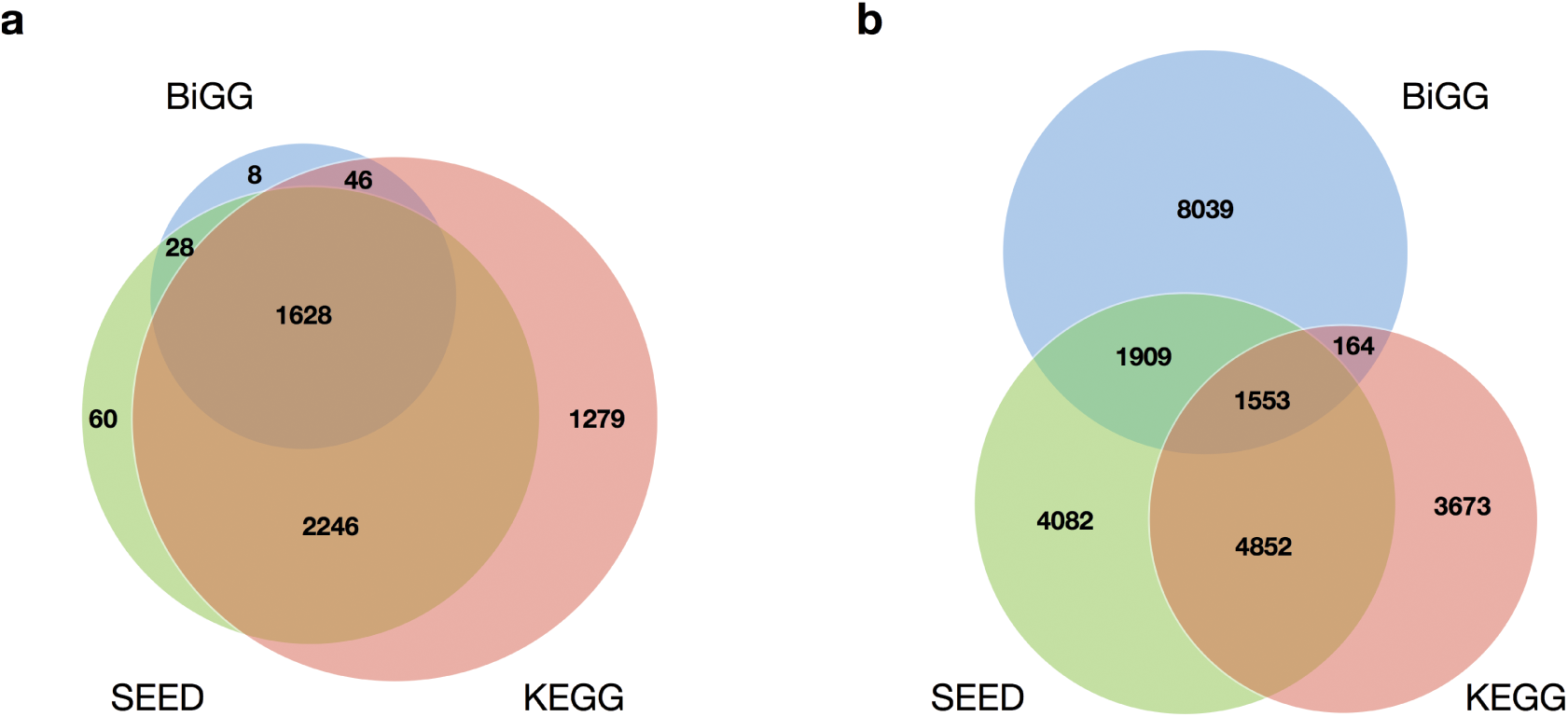
Comparison of the reaction coverage between different reaction databases: a) Total number of unique EC numbers present in each database; b) Total number of unique reactions present in each database. Data obtained from the MetaNetX database (version 3.0) [48].

## References

[1] M. A. Oberhardt, B. Ø. Palsson, J. A. Papin, Applications of genome-scale metabolic reconstructions, Molecular Systems Biology 5 (1) (2009) 320.

[2] A. Bordbar, J. M. Monk, Z. A. King, B. O. Palsson, Constraint-based models predict metabolic and associated cellular functions, Nature Reviews Genetics 15 (2) (2014) 107–120.

[3] D. Machado, M. J. Herrgård, Co-evolution of strain design methods based on flux balance and elementary mode analysis, Metabolic Engineering Communications 2 (2015) 85–92.

[4] A. Brochado, C. Matos, B. L. Møller, J. Hansen, U. H. Mortensen, K. Patil, Improved vanillin production in bakers yeast through in silico design, Microbial Cell Factories 9 (1) (2010) 84.

[5] T. Y. Kim, H. U. Kim, S. Y. Lee, Metabolite-centric approaches for the discovery of antibacterials using genome-scale metabolic networks, Metabolic Engineering 12 (2) (2010) 105–111.

[6] T. Shlomi, T. Benyamini, E. Gottlieb, R. Sharan, E. Ruppin, Genome-scale metabolic modeling elucidates the role of proliferative adaptation in causing the Warburg effect, PLoS Computational Biology 7 (3) (2011) e1002018.

[7] L. Väremo, I. Nookaew, J. Nielsen, Novel insights into obesity and diabetes through genome-scale metabolic modeling, Frontiers in Physiology 4 (2013) 92.

[8] A. Zelezniak, T. H. Pers, S. Soares, M. E. Patti, K. R. Patil, Metabolic Network Topology Reveals Transcriptional Regulatory Signatures of Type 2 Diabetes, PLoS Computational Biology 6 (4) (2010) e1000729.

[9] K. M. Havas, V. Milchevskaya, K. Radic, A. Alladin, E. Kafkia, M. Garcia, J. Stolte, B. Klaus, N. Rotmensz, T. J. Gibson, B. Burwinkel, A. Schneeweiss, G. Pruneri, K. R. Patil, R. Sotillo, M. Jechlinger, Metabolic shifts in residual breast cancer drive tumor recurrence, Journal of Clinical Investigation 127 (6) (2017) 2091–2105.

[10] A. R. Zomorrodi, C. D. Maranas, OptCom: a multi-level optimization framework for the metabolic modeling and analysis of microbial communities, PLoS Computational Biology 8 (2) (2012) e1002363.

[11] A. Zelezniak, S. Andrejev, O. Ponomarova, D. R. Mende, P. Bork, K. R. Patil, Metabolic dependencies drive species co-occurrence in diverse microbial communities, Proceedings of the National Academy of Sciences 112 (20) (2015) 6449–6454.

[12] O. Ponomarova, N. Gabrielli, D. C. Sévin, M. Mlleder, K. Zirngibl, K. Bulyha, S. Andrejev, E. Kafkia, A. Typas, U. Sauer, M. Ralser, K. R. Patil, Yeast Creates a Niche for Symbiotic Lactic Acid Bacteria through Nitrogen Overflow, Cell Systems 5 (4) (2017) 345–357e6.

[13] S. Freilich, R. Zarecki, O. Eilam, E. S. Segal, C. S. Henry, M. Kupiec, U. Gophna, R. Sharan, E. Ruppin, Competitive and cooperative metabolic interactions in bacterial communities, Nature Communications 2 (2011) 589.

[14] F. Salimi, K. Zhuang, R. Mahadevan, Genome-scale metabolic modeling of a clostridial co-culture for consolidated bioprocessing, Biotechnology Journal 5 (7) (2010) 726–738.

[15] N. Klitgord, D. Segre, Environments that Induce Synthetic Microbial Ecosystems, PLoS Computational Biology 6 (11) (2010) e1001002.

[16] M. Arumugam, J. Raes, E. Pelletier, D. L. Paslier, T. Yamada, D. R. Mende, G. R. Fernandes, J. Tap, T. Bruls, J.-M. Batto, et al., Enterotypes of the human gut microbiome, Nature 473 (7346) (2011) 174–180.

[17] S. Sunagawa, L. P. Coelho, S. Chaffron, J. R. Kultima, K. Labadie, G. Salazar, B. Djahan-schiri, G. Zeller, D. R. Mende, A. Alberti, et al., Structure and function of the global ocean microbiome, Science 348 (6237) (2015) 1261359.

[18] N. Fierer, Embracing the unknown: disentangling the complexities of the soil microbiome, Nature Reviews Microbiology 15 (10) (2017) 579–590.

[19] F. Meyer, D. Paarmann, M. D’Souza, R. Olson, E. M. Glass, M. Kubal, T. Paczian, A. Rodriguez, R. Stevens, A. Wilke, et al., The metagenomics RAST server–a public resource for the automatic phylogenetic and functional analysis of metagenomes, BMC Bioinformatics 9 (1) (2008) 386.

[20] C. S. Henry, M. DeJongh, A. A. Best, P. M. Frybarger, B. Linsay, R. L. Stevens, High-throughput generation, optimization and analysis of genome-scale metabolic models, Nature Biotechnology 28 (9) (2010) 977–982.

[21] R. Agren, L. Liu, S. Shoaie, W. Vongsangnak, I. Nookaew, J. Nielsen, The RAVEN toolbox and its use for generating a genome-scale metabolic model for Penicillium chrysogenum, PLoS Computational Biology 9 (3) (2013) e1002980.

[22] E. Pitkänen, P. Jouhten, J. Hou, M. F. Syed, P. Blomberg, J. Kludas, M. Oja, L. Holm, M. Penttilä, J. Rousu, et al., Comparative genome-scale reconstruction of gapless metabolic networks for present and ancestral species, PLoS Computational Biology 10 (2) (2014) e1003465.

[23] O. Dias, M. Rocha, E. C. Ferreira, I. Rocha, Reconstructing genome-scale metabolic models with merlin, Nucleic Acids Research (2015) gkv294.

[24] M. Kanehisa, S. Goto, KEGG: kyoto encyclopedia of genes and genomes, Nucleic Acids Research 28 (1) (2000) 27–30.

[25] I. Thiele, B. Ø. Palsson, A protocol for generating a high-quality genome-scale metabolic reconstruction, Nature Protocols 5 (1) (2010) 93–121.

[26] K. D. Pruitt, T. Tatusova, D. R. Maglott, NCBI reference sequences (RefSeq): a curated non-redundant sequence database of genomes, transcripts and proteins, Nucleic Acids Research 35 (suppl 1) (2007) D61–D65.

[27] J. D. Orth, T. M. Conrad, J. Na, J. A. Lerman, H. Nam, A. M. Feist, B. Ø. Palsson, A comprehensive genome-scale reconstruction of Escherichia coli metabolism2011, Molecular Systems Biology 7 (1) (2011) 535.

[28] Y.-K. Oh, B. O. Palsson, S. M. Park, C. H. Schilling, R. Mahadevan, Genome-scale reconstruction of metabolic network in Bacillus subtilis based on high-throughput phenotyping and gene essentiality data, Journal of Biological Chemistry 282 (39) (2007) 28791–28799.

[29] J. C. Xavier, K. R. Patil, I. Rocha, Integration of Biomass Formulations of Genome-Scale Metabolic Models with Experimental Data Reveals Universally Essential Cofactors in Prokaryotes, Metabolic Engineering 39 (2017) 200–208.

[30] M. B. Biggs, J. A. Papin, Managing uncertainty in metabolic network structure and improving predictions using EnsembleFBA, PLoS Computational Biology 13 (3) (2017) e1005413.

[31] A. B. Shreiner, J. Y. Kao, V. B. Young, The gut microbiome in health and in disease, Current Opinion in Gastroenterology 31 (1) (2015) 69.

[32] J. Aron-Wisnewsky, K. Clément, The gut microbiome diet, and links to cardiometabolic and chronic disorders, Nature Reviews Nephrology 12 (3) (2015) 169–181.

[33] M. G. Rooks, W. S. Garrett, Gut microbiota metabolites and host immunity, Nature Reviews Immunology 16 (6) (2016) 341–352.

[34] S. Magnúsdόttir, A. Heinken, L. Kutt, D. A. Ravcheev, E. Bauer, A. Noronha, K. Greenhalgh, C. Jäger, J. Baginska, P. Wilmes, et al., Generation of genome-scale metabolic reconstructions for 773 members of the human gut microbiota., Nature Biotechnology 35 (1) (2017) 81.

[35] J. Sung, S. Kim, J. Cabatbat, S. Jang, Y. Jin, G. Jung, N. Chia, P. Kim, Global metabolic interaction network of the human gut microbiota for context-specific community-scale analysis., Nature Communications 8 (2017) 15393.

[36] Z. A. King, J. Lu, A. Dräger, P. Miller, S. Federowicz, J. A. Lerman, A. Ebrahim, B. O. Palsson, N. E. Lewis, BiGG Models: A platform for integrating, standardizing and sharing genome-scale models, Nucleic Acids Research 44 (D1) (2016) D515–D522.

[37] R. Alcántara, K. B. Axelsen, A. Morgat, E. Belda, E. Coudert, A. Bridge, H. Cao, P. De Matos, M. Ennis, S. Turner, et al., Rheaa manually curated resource of biochemical reactions, Nucleic Acids Research 40 (D1) (2011) D754–D760.

[38] O. Dias, D. Gomes, P. Vilaça, J. Cardoso, M. Rocha, E. C. Ferreira, I. Rocha, Genome-Wide Semi-Automated Annotation of Transporter Systems, IEEE/ACM Transactions on Computational Biology and Bioinformatics (TCBB) 14 (2) (2017) 443–456.

[39] E. Noor, H. S. Haraldsdόttir, R. Milo, R. M. Fleming, Consistent estimation of Gibbs energy using component contributions, PLoS Computational Biology 9 (7) (2013) e1003098.

[40] J. O. Park, S. A. Rubin, Y.-F. Xu, D. Amador-Noguez, J. Fan, T. Shlomi, J. D. Rabinowitz, Metabolite concentrations, fluxes, and free energies imply efficient enzyme usage, Nature Chemical Biology 12 (7) (2016) 482.

[41] A. M. Feist, C. S. Henry, J. L. Reed, M. Krummenacker, A. R. Joyce, P. D. Karp, L. J. Broadbelt, V. Hatzimanikatis, B. Ø. Palsson, A genome-scale metabolic reconstruction for Escherichia coli K-12 MG1655 that accounts for 1260 ORFs and thermodynamic information, Molecular Systems Biology 3 (1) (2007) 121.

[42] B. Buchfink, C. Xie, D. H. Huson, Fast and sensitive protein alignment using DIAMOND, Nature Methods 12 (1) (2015) 59–60.

[43] J. Huerta-Cepas, K. Forslund, L. P. Coelho, D. Szklarczyk, L. J. Jensen, C. von Mering, P. Bork, Fast genome-wide functional annotation through orthology assignment by eggNOG-mapper, Molecular Biology and Evolution msx148.

[44] A. R. Wattam, D. Abraham, O. Dalay, T. L. Disz, T. Driscoll, J. L. Gabbard, J. J. Gillespie, R. Gough, D. Hix, R. Kenyon, et al., Patric, the bacterial bioinformatics database and analysis resource, Nucleic acids research 42 (D1) (2013) D581–D591.

[45] D. Machado, S. Andrejev, framed 0.4.0 (2017). doi:10.5281/zenodo.1048261. URL https://doi.org/10.5281/zenodo.1048261

[46] B. J. Bornstein, S. M. Keating, A. Jouraku, M. Hucka, LibSBML: an API library for SBML, Bioinformatics 24 (6) (2008) 880–881.

[47] T. Yamada, I. Letunic, S. Okuda, M. Kanehisa, P. Bork, iPath2. 0: interactive pathway explorer, Nucleic acids research 39 (suppl 2) (2011) W412–W415.

[48] S. Moretti, O. Martin, T. V. D. Tran, A. Bridge, A. Morgat, M. Pagni, MetaNetX/MNXref – reconciliation of metabolites and biochemical reactions to bring together genome-scale metabolic networks, Nucleic Acids Research 44 (D1) (2015) D523–D526.

